# High spatiotemporal resolution radial encoding single vessel fMRI

**DOI:** 10.1101/2023.12.26.573112

**Authors:** Yuanyuan Jiang, Patricia Pais-Roldán, Rolf Pohmann, Xin Yu

## Abstract

High-field preclinical functional MRI (fMRI) has enabled the high spatial resolution mapping of vessel-specific hemodynamic responses, i.e. single-vessel fMRI. In contrast to investigating neuronal sources of the fMRI signal, single vessel fMRI focuses on elucidating its vascular origin, which can be readily implemented to identify vascular changes relevant to vascular dementia or cognitive impairment. However, the limited spatial and temporal resolution of fMRI has hindered hemodynamic mapping of intracortical microvessels. Here, we implemented the radial encoding MRI scheme to measure BOLD signals of individual vessels penetrating the rat somatosensory cortex. Radial encoding MRI is employed to map cortical activation with a focal field of view (FOV), allowing vessel-specific functional mapping with 50×50 µm in-plane resolution at 1 to 2 Hz sampling rate. Besides detecting refined hemodynamic responses of intracortical micro-venules, the radial encoding-based single-vessel fMRI enables the differentiation of the intravascular and extravascular effects from the draining venules.

## Introduction

Conventional functional magnetic resonance imaging (fMRI) methods^1–4^ are developed to measure hemodynamic response as the surrogate of neuronal activity. The vascular origin of fMRI signals can be specified as changes in blood volume, flow, and oxygenation saturation driven by neurovascular coupling. Since the exact volume contribution of the cerebrovasculature to the brain is less than ∼2-4%^5–7^, fMRI signals of a given voxel with sub-millimeter-to-millimeter cubic size are considered to present brain function in a relatively macroscopic scale. Although there is no consensus to treat voxel-wise fMRI signals as populational coding of cellular-specific neuronal activity, the emerging optogenetic tools in combination with fMRI have opened a new avenue to decipher cellular component contribution to the fMRI signal in animal models^8–10^. However, to provide an insightful interpretation of the fMRI signal, the contribution of vascular components to the fMRI signal should be better elucidated, particularly with the evolved MR technology to improve the spatiotemporal resolution of fMRI brain mapping.

With state-of-the-art high-field MR technology, human brain fMRI has acquired functional maps with a spatial resolution of 300-500 µm voxels^11–15^, which could well separate the pial vessels, with near hundred-micron diameters. In contrast to human brain mapping, the high-field rodent fMRI has enabled 2D slice image acquisition with 100×100 µm in-plane resolution across the cortex^16^. Because the rodent brain has a smooth surface with micro-vessels radially distributed throughout the cortex, the vascular partial volume contribution to the high-resolution fMRI signals of rodent brains is not negligible^17^. Given the T2* extravascular amplification effect, the “single-vessel” fMRI enables the detection of arteriole-dominated cerebral blood volume (CBV) and venule-dominated blood-oxygen-level-dependent (BOLD) fMRI signals from intracortical vessel voxels^9, 16, 18, 19^. A recent study has also detected vessel-specific functional cerebral blood volume (CBF) velocity changes with high-resolution phase-contrast MRI^20^. However, the increased spatial and temporal resolution of single-vessel fMRI acquisition leads to an inevitable signal-to-noise ratio (SNR) loss. Previous single-vessel fMRI methods apply reshuffled k-t space fast low angle shot (FLASH) or balanced steady-state free precession (bSSFP) sequences with focal field-of-view (FOV) to ensure sufficient SNR while maintaining the high spatiotemporal resolution^16, 21, 22^. These methods also acquire single-vessel images with less distorted images than the conventional echo-planar imaging (EPI) method, but it remains challenging to push the resolution higher given the interdependent spatial, temporal resolution and SNR.

To achieve the finest spatial scale for single-vessel fMRI, we implement the radial encoding MRI scheme to measure the individual vessels penetrating the rat somatosensory cortex. In contrast to the cartesian acquisition schemes, the radial encoding offers continuous updating of the center of the k-space and pushes the 50×50 µm spatial resolution with a 1 to 2 Hz sampling rate by defining the arbitrary number of projections in the azimuthal direction. Besides detecting refined hemodynamic maps of intracortical vessels, the radial encoding based single-vessel fMRI offers the opportunity to distinguish the intravascular and extravascular effects from the cortical vessels.

## Results

We implemented the high-resolution radial encoding based single-vessel fMRI acquisition for *in vivo* hemodynamic measurement of individual penetrating arterioles and venules of anesthetized rats with a 14 T MR scanner. To better align the 2D radial encoding slice (FOV 9.6×9.6 mm^2^) along the cortical region of interest (**Supplementary Fig 1**), we have adjusted the animal holder with a turning ring to allow one more degree of freedom to adjust the head rotation along the z-axis of the MRI scanner.

### Varied FOV acquisition for single-vessel radial encoding fMRI

Since radial encoding applies the frequency-encoding projection in a radial scheme, the aliasing effect of phase encoded direction will not affect the image acquisition. We first acquired the single-vessel radial encoding fMRI images with three FOVs: 9.6×9.6, 6.4×6.4, and 4.8×4.8 mm². **Fig 1A** shows the overlaid BOLD fMRI maps on the multi-gradient echo (MGE)-based anatomical arteriole-venule (A-V) map, highlighting the activated venule voxels from the primary forepaw somatosensory cortex (FP-S1) of anesthetized rats. As previously reported^9, 23^, we applied an MGE sequence to acquire the A-V map, where the venule voxels appear as dark dots due to fast T2* decay, while the arteriole voxels remain bright dots due to the in-flow effect. The peak BOLD signals were mainly located on venule voxels, indicating venule-dominated BOLD responses. Under the block-design paradigm (stimulation on/off epochs), robust positive BOLD fMRI signals can be measured from individual venules with varied FOV. The reduction in FOV not only preserved the active cortical region but also allowed to decrease the time of repetition (TR) from 1.5 s to 0.5 s. The averaged time course of voxels from a single venule exhibited a robust response, and the overall contrast-to-noise ratio was high enough to detect single venule-specific BOLD responses in the faster sampling scheme (**Fig 1B&C**). **Fig 1D** shows an example of the A-V map with highlighted venule and arteriole ROIs. The BOLD fMRI signals of venule ROIs were significantly higher than those of arteriole ROIs, which were detected at 4s and 15s duration stimulation paradigms with different TRs (0.5s, 1s, 1.5s) (**Fig 1E&F** & **Supplementary Fig 2**). These findings indicate the potential advantages of using smaller FOVs to detect vessel-specific hemodynamic responses with a faster sampling rate using single-vessel radial encoding fMRI.

**Figure 1.**
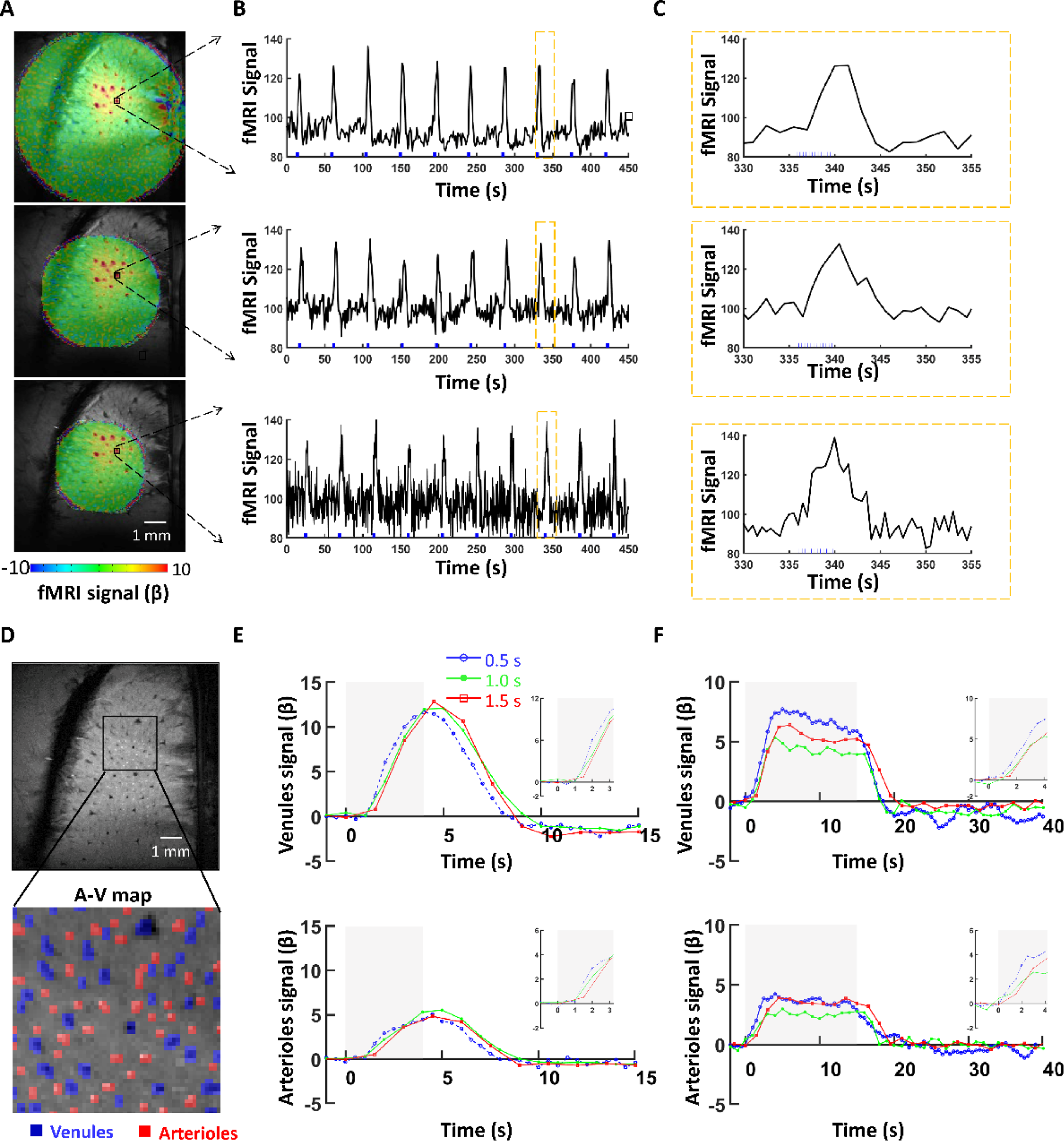
The single-vessel radial encoding fMRI acquisition to achieve arbitrary FOV. (**A**) By increasing the radial encoding projections, the FOV was reduced from 9.6×9.6 mm^2^ to 6.4×6.4 mm^2^ and 4.8×4.8 mm^2^ while maintaining covering the main responsive somatosensory cortex (FP-S1) cortex. (**B**) The BOLD fMRI signal from a single venule in each FOV acquisition. (**C**) The zoom-in view of a faster sampling scheme from a single vessel captures real-time BOLD responses. (**D**) The vessel-dominated BOLD fMRI A-V map from the 2D MGE acquisition. The arteriole voxels were marked as red ROIs, and the venule voxels as blue ROIs. (**E**) The averaged venules and arterioles of the evoked BOLD fMRI for different FOV acquisitions with sampling rates (0.5, 1, and 1.5 s). The block-design paradigm for the forepaw stimulation train (330 μs, 1.5 mA) was delivered at 3 Hz for 4 s (**E**) and 3 Hz for 15 s (**F**). The stimulation period is shown as a light-gray shaded area. The zoomed views of the onset are outlined in each upper panel.

### Varied spatial resolution for single-vessel radial encoding fMRI

By increasing the azimuthal projections at the same FOV (9.6×9.6 mm²) from 75 to 100 and to 150 projections, single-vessel radial encoding fMRI enabled to increase the in-plane resolution of hemodynamic mapping from100×100 µm^2^ to 75×75 µm^2^ and 50×50 µm^2^, respectively. When comparing the anatomical characterization of vessel voxels from different raw images, the most distinct and well-defined spreading function of individual venules could be identified in radial encoding images of 50×50 µm^2^ (**Fig 2A**). BOLD functional maps were also obtained by single-vessel radial encoding fMRI with different spatial resolutions, demonstrating much more refined hemodynamic maps overlaying on the A-V maps (**Fig 2B**). We further analyzed the hemodynamic characteristics of vessel-specific BOLD fMRI signals at different spatial resolutions. Using the A-V map-based vessel ROIs, the venule-specific BOLD responses of 50 µm resolution represent relatively lower amplitude than the 100 µm resolution due to altered partial volume extravascular effect from different voxel sizes (**Fig 2C**). Meanwhile, the BOLD signal from individual venules ROIs acquired at 50 µm resolution shows sharper profile distribution and higher peak responses than 75 µm under the same TE acquisition (**Fig 2D, E**), indicating that more refined vessel-specific BOLD responses are detected with the version of the sequence employing higher spatial resolution.

**Figure 2.**
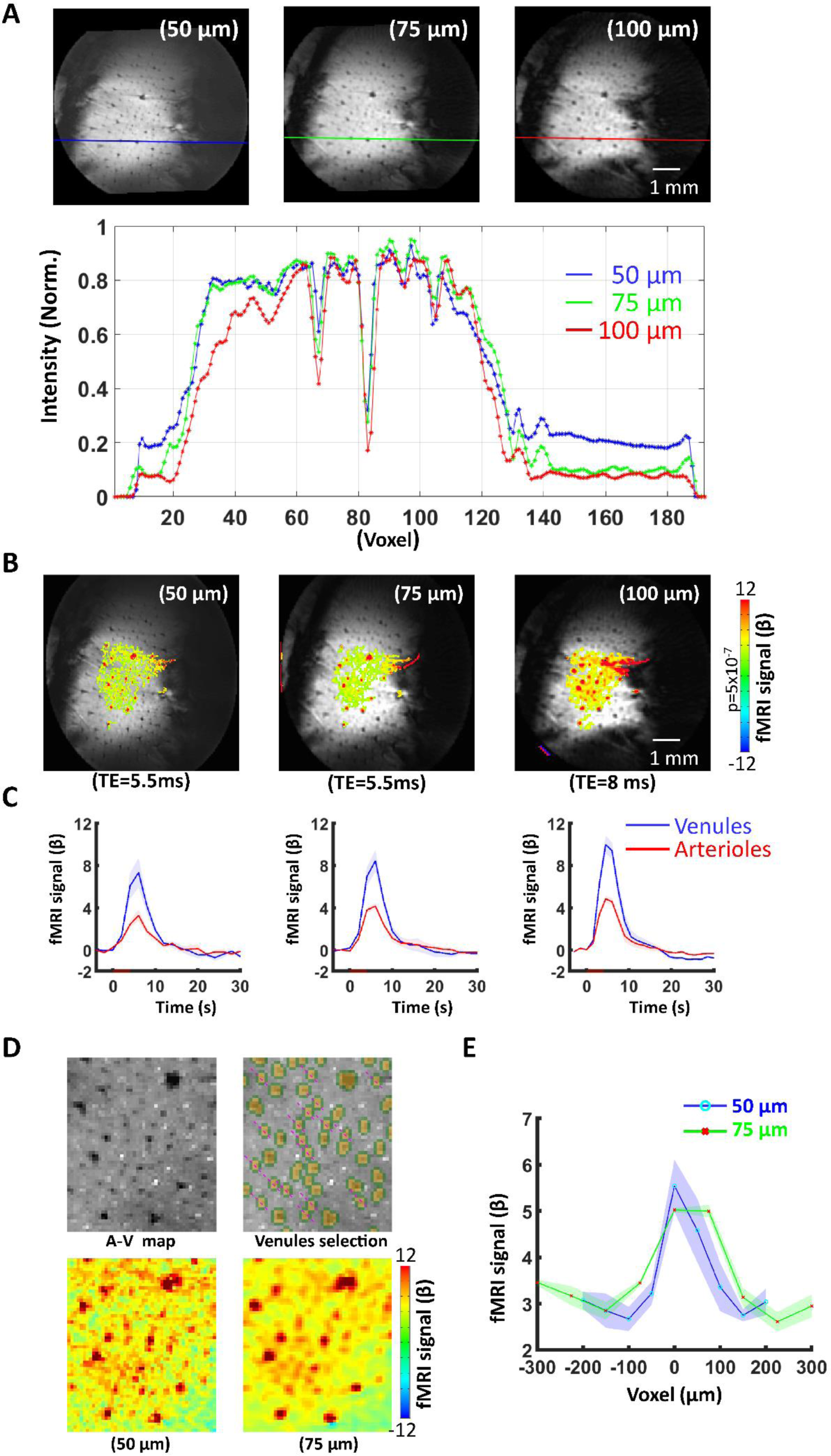
High-resolution single-vessel radial encoding fMRI recording acquisition. (**A**) The BOLD fMRI signal from a single venule in each FOV acquisition and corresponding line intensity profile across the vessels. A clear point-spreading function of the individual vessel can be indemnified from the 50μm resolution fMRI. (**B**) The BOLD fMRI maps at various resolutions (50 µm and 75 µm with TE=5 ms, and 100 µm with TE=8 ms) were superimposed within the same 9.6 mm × 9.6 mm FOV. (**C**) The BOLD fMRI signal originating from venules and arterioles in the active cortex region of different resolution acquisition (n=4 rats, mean± SD). (**D**) The parenchyma voxel profile extraction for voxels within the individual venules mask (one venule voxel) and the corresponding BOLD fMRI distribution at 50 µm and 75 µm resolutions under the same TE (5 ms). (**E**) The averaged venules-specific fMRI profile patterns at 50 μm and 75 μm resolution from D (n=4 rats, mean± SEM).

Moreover, the single-vessel radial encoding fMRI maps obtained at different resolutions enabled the direct characterization of distinct intravascular and extravascular effects from large pial vessels. Depending on the slice localization, the low spatial resolution scheme (100×100 µm) showed not only positive BOLD signals in venule voxels but also negative BOLD signals in voxels covering the pial surface vein (**Fig 3A, Supplementary Fig 3)**. The negative BOLD signal can be caused by the passive venule dilation (i.e. reduced T2* signal due to the CBV effect)^24–26^, which will mostly affect voxels with 100×100 µm in-plane size, covering both blood and parenchymal tissue. The surrounding positive BOLD signals are caused by the typical extravascular effect of oxy/deoxy-hemoglobin ratio increase. However, the negative signal disappeared at higher spatial resolution radial encoding fMRI mapping with positive BOLD signals surrounding the pial veins (**Fig 3B** &**Supplementary Fig 4**). This is possibly due to the fact that a smaller voxel size enables the detection of the intravascular blood signals that are diminished at high magnetic field^27^ without the negative CBV effect (**Fig 3C)**. This radial encoding based high-resolution fMRI has presented highly refined vessel-specific hemodynamic responses of rat brains, enabling the dissection of vessel-specific contributions to fMRI signals.

**Figure 3.**
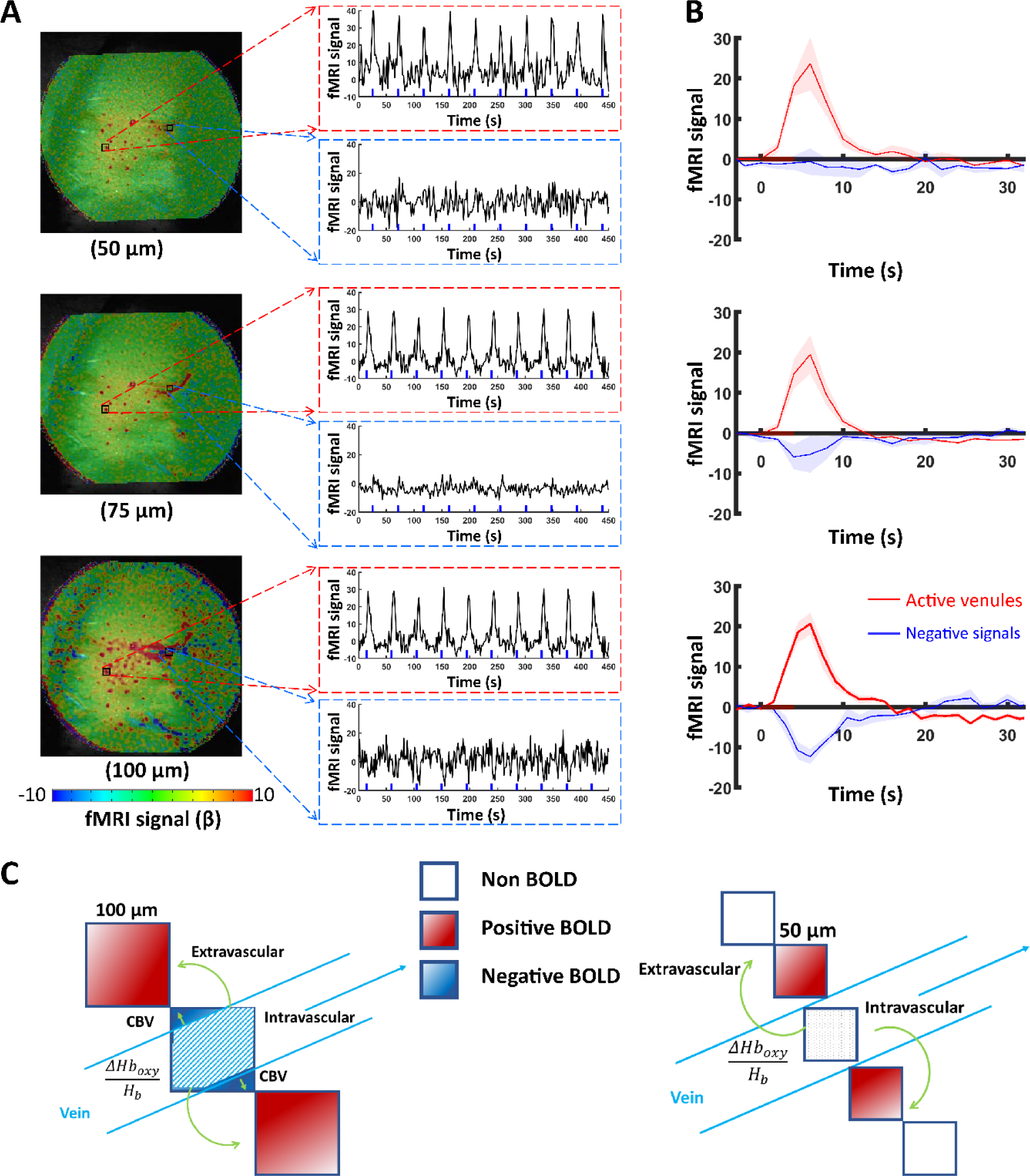
Intravascular effect of draining veins of high-resolution single-vessel radial encoding fMRI. (**A**) The distributions of BOLD fMRI maps in-plane resolution range from 50 to 100 μm (FOV 9.6 × 9.6 mm^2^). The positive fMRI signals from single venules and negative signals surrounding the pial veins. The position and voxel signal time courses in active venules and voxels surrounding the draining veins (**Supplementary Figure 3&4**). (**B**) The average positive and negative BOLD signals from different animals (n=4 rats, mean± SEM). (**C**) The schematic diagram illustrates the impact on intravascular and extravascular effects caused by changes in voxel size.

### Comparison of single-vessel radial encoding and bSSFP fMRI

We also conducted a comparison between single-vessel radial encoding and bSSFP fMRI focusing on the 100×100 µm in-plane resolution in the same animals. Both methods revealed highly robust positive BOLD fMRI signals located at the venule voxels (**Fig 4A**), demonstrating the compelling consistency of the single-vessel venule-dominated BOLD detection (**Fig 4B**). Meanwhile, it is important to note the presence of potential bSSFP banding artifacts in the cortex due to the field inhomogeneity (**Fig 4C**). **Supplementary movie 1** presents representative evidence of these artifacts, and the pattern of banding artifacts evolved during the acquisition due to the altered gradient/shimming coil temperature. Also, the radial encoding fMRI dataset showed significantly higher tSNR than bSSFP datasets (**Fig 4C-E**), indicating better detectability of the single-vessel radial encoding to measure the vessel-specific hemodynamic responses.

**Figure 4.**
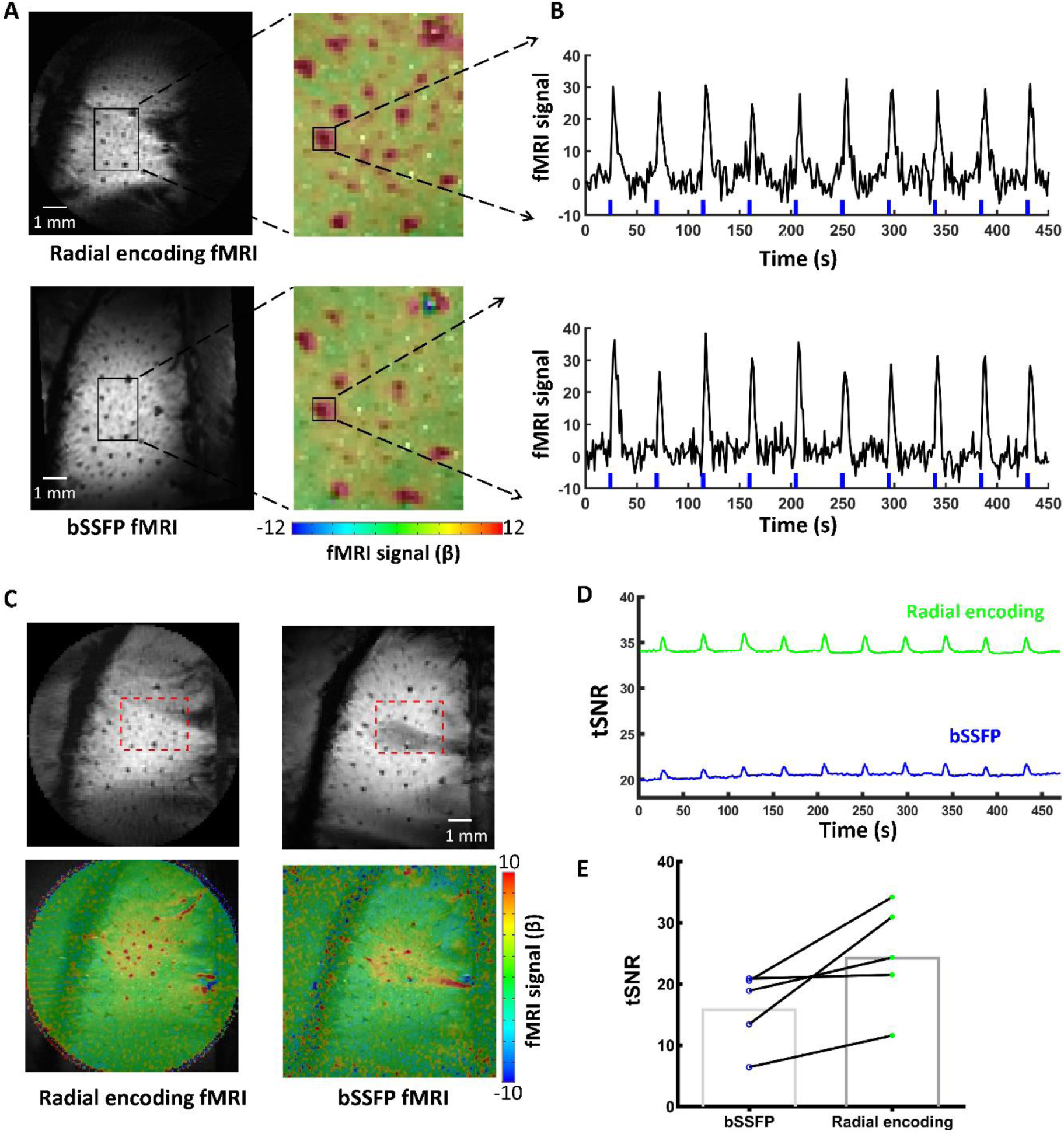
Comparison of single-vessel radial encoding fMRI and bSSFP fMRI. (**A**) The 100 μm in-plane resolution single-vessel radial encoding fMRI and bSSFP fMRI image and BOLD maps (FOV 9.6 × 9.6 mm^2^, 40% transparency fMRI map). **(B**) The positive fMRI signals from single venules from single-vessel radial encoding fMRI and bSSFP fMRI. (**C**) The representative images and fMRI maps of single-vessel radial encoding and bSSFP acquisition. In bSSFP acquisition, banding artifacts are evident in the active cortex region of some animals, leading to a reduction in fMRI signals. (**D**) The representative time course of temporal Signal-to-Noise Ratio (tSNR) of single-vessel radial encoding and bSSFP fMRI acquisition. The rectangular mask for the active region of interest (ROI) is displayed in panel C. (**E)** A paired comparison of the mean tSNR values for single-vessel bSSFP and radial encoding fMRI from the same animal acquisition (n=5 rats, paired t-test, p<0.02).

## Methods

### Animal preparation

All animal surgical and experimental procedures were in full compliance with the guide for the care and use of laboratory animals and approved by the Massachusetts General Hospital Institutional Animal Care and Use Committee. Male Sprague Dawley rats (∼250g) were intubated with a mechanical ventilator (SAR-830, CWE, USA) and anesthetized with 2% isoflurane. Blood pressure monitoring and anesthetics infusion took place via femoral artery and vein catheterization. The isoflurane was discontinued after the i.v. bolus injection of α-chloralose (Sigma-Aldrich, 80mg/kg) through the femoral vein. During MR acquisition, the α-chloralose (26.5mg/kg/h) mixed with pancuronium (Zemuron, 2 mg/kg/h) was infused to immobilize the rats. Also, a heating pad was used to maintain the rectal temperature of rats at ∼37°C for the duration of the experiment. All relevant physiological parameters were constantly monitored during scanning: heart rate, arterial blood pressure, the pressure of the tidal ventilation (Biopac MP 160, Biopac Systems, USA), and end-tidal CO_2_ (Capnometer, Novametrix).

### MRI acquisition

MRI images were acquired using a 14 T, 26 cm horizontal bore magnet (Magnex Scientific) interfaced through the Bruker Advance III console (Bruker Corporation). A 12 cm magnet gradient set was equipped with a strength of 100 G/cm and a 150 μs rise time (Resonance Research Inc.) in the scanner. The stimulation paradigm was triggered directly through the MRI scanner and was controlled by Master-9 A.M.P.I system (Jerusalem, Israel). The triggering pulses from the MRI scanner were also recorded by a Biopac system (MP160, Biopac Systems, USA). To map the sensory-evoked single-vessel fMRI, a pair of stimulation electrodes were placed on the forepaw to deliver trains of 1.5 mA pulses (300 μs) at 3Hz for 4s or at 3Hz for 15s in each fMRI epoch. A 6 mm transceiver coil was constructed and attached to the rat skull covering the somatosensory cortex.

### Single-vessel multi-gradient echo (MGE) imaging

To acquire the anatomical arteriole-venule (A-V) map, a 2D multiple gradient echo (MGE) sequence was implemented with the following parameters: TE: 2.5, 5, 7.5, 10, 12.5, 15 ms; TR: 50 ms; slice thickness: 500 μm; flip angle: 55°; matrix: 192×192; in-plane resolution: 50×50 μm. The single vessel map is acquired by averaging the MGE images acquired from the second echo to the fourth echo.

### Single-vessel radial encoding BOLD fMRI

We employed radial encoding acquisition with varying parameters to investigate spatial and temporal freedom in fMRI. For temporal freedom acquisition, the slice thickness 500 µm an in-plane resolution of 100×100 µm, flip angle of 15° were utilized. The following parameters were used for a FOV of 9.6×9.6 mm^2^ acquisition: TE of 8 ms; TR of 19.738 ms; 76 projections; matrix size of 96×96. For a reduced FOV of 6.4×6.4 mm^2^: TE of 8 ms; TR of 20 ms; 50 projections, matrix size of 64×64. To achieve a sampling rate of 2 Hz and cover the responsive cortical region with a FOV of 4.8×4.8 mm^2^, we applied TE of 8 ms and TR of 13.159 ms; 38 projections; matrix size of 64×64. For FOV 9.6×9.6 mm^2^, the block design was 25 pre-stimulation scans, 1 scan during stimulation, and 29 post-stimulation scans with 20 epochs for each trial and a total scan duration of 15 minutes and 37 seconds. For FOV 6.4×6.4 mm, the block design was 25 pre-stimulation scans, 1 scan during stimulation, and 44 post-stimulation scans with 20 epochs for each trial with a total scan duration of 15 minutes and 25 seconds. For FOV 4.8×4.8 mm, the block design was 50 pre-stimulation scans, 1 scan during stimulation, and 89 post-stimulation scans with 20 epochs for each trial with a total scan duration of 15 minutes and 25 seconds.

For investigating different in-plane spatial resolutions of 50, 75, and 100 µm, we recorded single-vessel fMRI images with varying numbers of projections and matrix sizes. Specifically, for 50 µm resolution, we used a matrix size of 192×192 and 150 projections. For 75 µm resolution, the matrix size was 128×128, and we used 100 projections. For 100 µm resolution, the matrix size was 96×96, and we used 75 projections. These settings allowed us to assess the impact of spatial and temporal freedom acquisition in radial encoding fMRI. For the in-plane resolution of 50 and 75 µm acquisition, the block design was 25 pre-stimulation scans, 1 scan during stimulation, and 22 post-stimulation scans with 20 epochs for each trial with a total scan duration of 16 minutes and 10 seconds.

### Balanced steady-state free precession (bSSFP) single-vessel BOLD fMRI

For real-time bSSFP fMRI acquisition, the following parameters were used: TE of 7.8 ms; TR of 15.625 ms; flip angle of 15°; FOV of 9.6×9.6 mm^2^; matrix size of 96×96; one slice repetition time of 1.5 s. The fMRI block design consisted of 25 pre-stimulation scans, 1 scan during stimulation, and 44 post-stimulation scans with 20 epochs. The total acquisition duration of each trial was 15 minutes and 37 seconds. The geometry of bSSFP was set to the same as the 100 µm radial encoding MRI.

### Data analysis

All data preprocessing and analysis were performed using the software package Analysis of Functional NeuroImages (AFNI) (NIH, Bethesda, USA) and MATLAB (MathWorks, Natick, USA). All relevant fMRI analysis source codes can be downloaded from (https://afni.nimh.nih.gov/). A detailed description of the processing procedure can be found in a previous study^9, 20^. To register the evoked single-vessel fMRI images with the 2D anatomical A-V map, the tag-based registration method was applied. The classification of individual vessel voxels was based on their signal intensity. Voxels with signal intensity higher than the mean signal intensity plus three times the standard deviation were color-coded in red, representing arterioles. On the other hand, voxels with signal intensity lower than the mean signal intensity minus three times the standard deviation of local areas (in a 5×5 kernel) were color-coded in blue, representing venules. Multiple trials of block-design time courses were averaged for each animal. No additional smoothing step was applied. 3dDeconvolve module developed in AFNI was used to map hemodynamic response based on the “block function”. The hemodynamic model is defined as follows:

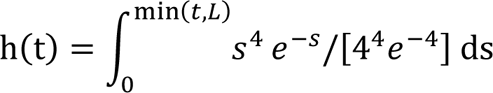

L was the duration of the response. BLOCK (L,1) is a convolution of a square wave of duration L with gamma variate function = s^4^e^-s^/4^4^e^-4^, making a peak amplitude of block response = 1. The fMRI β-value was calculated to measure the amplitude of the fMRI responses at each TR. tSNR were computed from the mean value of 30×20 voxels in the center of the activation region across all the fMRI time points, divided by the temporal standard deviation of the voxels. For the group analysis, a Student’s t-test was performed, and error bars in all figures represent the mean ± standard error of the mean (SEM). The p values < 0.05 were considered statistically significant.

## Discussion

High spatial resolution fMRI in animal models has evolved to identify individual penetrating microvessels with distinct vessel-specific hemodynamic responses. This single-vessel fMRI approach allows non-invasive detection of large-scale hemodynamic signals from large animals without spectral-specific signal attenuation through the skull, e.g. ultrasound or optoacoustic imaging^28–30^. The present study has implemented the radial encoding scheme to further push the spatial resolution of fMRI to 50 µm in-plane with sub-second TR, aiming to dissect the vascular contributions to fMRI signals.

### Advantage of the radial encoding scheme for ultra-high resolution fMRI

With the conventional cartesian trajectory k-space acquisition, faster sampling can be achieved by using a reduced matrix and phase encoding steps with a smaller in-plane FOV^31^, but the aliasing effect can lead to fold-over artifacts. Anti-aliasing encoding schemes can be used but this often comes at the cost of decreased temporal resolution. Although the echo planar imaging (EPI) method enables fast sampling through the echo trains with a prioritized wide range of brain coverage, the detected hemodynamic response can suffer from distortion due to the embedded T2* effect during acquisition. To enable the acquisition of small FOV without aliasing artifacts, the radial encoding sampling scheme has been inmplemented^32, 33^. This radial encoding method presents numerous advantageous features for ultra-high resolution fMRI brain mapping. To mitigate the aliasing artifacts, radial encoding implemented a radial frequency encoding scheme without the fold-over issues due to the phase-encoding scheme. Also, radial encoding enables repeated sampling in the k-space center to ensure an overall uniform contrast. Interestingly, by under-sampling in the azimuthal direction, image acquisition can be accelerated to only focus on a smaller FOV (e.g. 4.8×4.8 mm^2^) with high spatial resolution, enabling the single-vessel fMRI with a faster sampling rate. This approach is advantageous because it focuses on the region of interest where the most signal energy is frequently concentrated, allowing for high temporal resolution. A constant azimuthal profile spacing (111.246°) based on the golden ratio can further be applied to optimize image reconstruction from a more uniform profile distribution in radial MRI^33–35^. By prioritizing the ROI and capturing the most relevant signal information, the single-vessel GRE method enhances the accuracy and fidelity of the hemodynamic response measurement.

### Comparison to the previously developed single-vessel fMRI methods

The usage of reshuffled k-t space FLASH for single-vessel fMRI has demonstrated remarkable capabilities in achieving high spatiotemporal resolution^9^, however, it is not a real-time fMRI acquisition. Recently, a new approach called direct imaging of neuronal activity (DIANA)^36^, based on 2D shuffled line scanning has been proposed^9, 37, 38^. There has been an ongoing debate regarding the reproducibility of the DIANA method^22–24^. One ongoing concern is the temporal aliasing issue with this reshuffled k-t space trajectory^39–41^, which could make the detected ultra-fast dynamic signals confounded by both physiological and non-physiological noises.

An alternative method, the bSSFP method has gained recognition for its ability to offer real-time monitoring of hemodynamic signals and superior SNR efficiency compared to other pulse sequences^21, 22^. Previously, He et al have applied the bSSFP for single-vessel hemodynamic signal detection with an in-plane resolution of 100 µm^16, 39^. Typically, SSFP applications focus on the pass-band region, where a high SNR per unit time can be attained while being relatively less affected by precise off-resonance frequencies. To mitigate this off-resonance banding artifact, careful shimming adjustments need to be periodically applied during the experiment. As long as effective shimming is employed to confine the frequency range within the pass-band in the anatomical region of interest, the resulting image will exhibit uniform signal and contrast. It is worth noting that when using high spatial resolution from focal FOV with short TR/TE, the demanded strong gradient in the high-duty cycle bSSFP sequence leads to localized temperature increases in the gradient to alter the shimming outcomes during scanning (**Fig 4A**). This challenging issue is a main course of the banding artifacts of the bSSFP-based single-vessel fMRI.

The radial encoding scheme allows the reconstruction of the fMRI images based on the number the azimuthal projections. This method provides the flexibility to alter the FOV, as well as the corresponding spatial and temporal resolution. One challenging issue is to position the brain region of interest to the geometrical center of the magnet (or the gradient). We have produced an animal holder to ease the animal head positioning inside the MRI scanner (**Supplementary Fig 1**), allowing geometrical orientation setup for radial encoding fMRI to target the region of interest of rat brains.

### The focus shifted from functional MRI to hemodynamic MRI

Numerous efforts have been made to improve the spatial specificity of fMRI by eliminating the draining vein-mediated mislocalization of functional maps^5, 27, 42–47^. These studies aim to present the brain function with fMRI by emphasizing the hemodynamic responses from intracortical voxels enriched with arterioles and capillaries, which have a closer proximity to the neuronal sources^48–50^. One fundamental difference between the high-resolution single-vessel fMRI with the previous studies is that it targets the vascular origin of fMRI. Thus, in contrast to the “functional” term of fMRI, the single-vessel fMRI approach focuses on the hemodynamic features, which makes it more appropriate to be called a single-vessel “hemodynamic” (h)MRI.

Different from the advanced optical imaging, optoacoustic, and functional ultrasound imaging schemes, the unique non-invasive and global mapping features of single-vessel fMRI ensures the detection of less disturbed neurovascular coupling, as well as being less dependent on the vascular flow orientation than the Doppler effect. Nevertheless, given the shifted focus on vascular specificity, the challenge of single-vessel fMRI is to detect more refined micro-vessels with high spatiotemporal resolution. Single-vessel radial encoding fMRI proposed in this study is an ongoing effort to map the penetrating micro-vessels from rodent brains. In particular, the 50×50 µm in-plane resolution single-vessel maps enables the well-characterized BOLD fMRI signals from refined arteriole and venule voxels, attributing more vessel-specific hemodynamic features.

### Summary

Single-vessel radial encoding fMRI involves aligning the k-space line radially to increase temporal resolution when mapping focal cortical regions with reduced artifacts. The ability to assign different projection numbers provides unique flexibility to develop more efficient and customizable data acquisition strategies, resulting in improved spatial resolution and localization specificity in fMRI studies. Moreover, this dynamic imaging approach demonstrated superior tSNR compared to bSSFP fMRI, making it particularly suitable for real-time fMRI applications. This radial encoding scheme enables dynamic imaging using continuous data acquisition and retrospective reconstruction of image series, which can be combined with respiratory or cardiac motion and compressed sensing reconstruction techniques^33, 51^. This combination further benefits real-time single-vessel BOLD fMRI, CBV, and cerebral blood flow studies, making it a valuable tool for advanced brain functional mapping with high-field MRI scanners.

## References

1. Kwong KK, Belliveau JW, Chesler DA, Goldberg IE, Weisskoff RM, Poncelet BP, Kennedy DN, Hoppel BE, Cohen MS, Turner R, et al. Dynamic magnetic resonance imaging of human brain activity during primary sensory stimulation. Proceedings of the National Academy of Sciences of the United States of America. 1992;89(12):5675–9. PubMed PMID: 1608978; PMCID: PMC49355.

2. Bandettini PA, Wong EC, Hinks RS, Tikofsky RS, Hyde JS. Time course EPI of human brain function during task activation. Magn Reson Med. 1992;25(2):390–7. Epub 1992/06/01. doi: 10.1002/mrm.1910250220. PubMed PMID: 1614324.

3. Ogawa S, Tank DW, Menon R, Ellermann JM, Kim SG, Merkle H, Ugurbil K. Intrinsic signal changes accompanying sensory stimulation: functional brain mapping with magnetic resonance imaging. Proceedings of the National Academy of Sciences of the United States of America. 1992;89(13):5951–5. PubMed PMID: 1631079; PMCID: PMC402116.

4. Belliveau JW, Kennedy DN, Jr., McKinstry RC, Buchbinder BR, Weisskoff RM, Cohen MS, Vevea JM, Brady TJ, Rosen BR. Functional mapping of the human visual cortex by magnetic resonance imaging. Science. 1991;254(5032):716–9. Epub 1991/11/11. doi: 10.1126/science.1948051. PubMed PMID: 1948051.

5. Kim S-G, Ogawa S. Biophysical and Physiological Origins of Blood Oxygenation Level-Dependent fMRI Signals. Journal of Cerebral Blood Flow & Metabolism. 2012;32(7):1188–206. doi: 10.1038/jcbfm.2012.23. PubMed PMID: 22395207.

6. Weber B, Keller AL, Reichold J, Logothetis NK. The Microvascular System of the Striate and Extrastriate Visual Cortex of the Macaque. Cerebral Cortex. 2008;18(10):2318–30. doi: 10.1093/cercor/bhm259.

7. Ji X, Ferreira T, Friedman B, Liu R, Liechty H, Bas E, Chandrashekar J, Kleinfeld D. Brain microvasculature has a common topology with local differences in geometry that match metabolic load. Neuron. 2021;109(7):1168–87.e13. doi: 10.1016/j.neuron.2021.02.006.

8. Lee JH, Durand R, Gradinaru V, Zhang F, Goshen I, Kim DS, Fenno LE, Ramakrishnan C, Deisseroth K. Global and local fMRI signals driven by neurons defined optogenetically by type and wiring. Nature. 2010;465(7299):788–92. doi: 10.1038/nature09108. PubMed PMID: 20473285; PMCID: PMC3177305.

9. Yu X, He Y, Wang M, Merkle H, Dodd SJ, Silva AC, Koretsky AP. Sensory and optogenetically driven single-vessel fMRI. Nat Methods. 2016;13(4):337–40. doi: 10.1038/nmeth.3765. PubMed PMID: 26855362.

10. Schmid F, Wachsmuth L, Schwalm M, Prouvot P-H, Jubal ER, Fois C, Pramanik G, Zimmer C, Faber C, Stroh A. Assessing sensory versus optogenetic network activation by combining (o)fMRI with optical Ca2+ recordings. Journal of Cerebral Blood Flow & Metabolism. 2015;36(11):1885–900. doi: 10.1177/0271678X15619428.

11. Feinberg DA, Vu AT, Beckett A. Pushing the limits of ultra-high resolution human brain imaging with SMS-EPI demonstrated for columnar level fMRI. NeuroImage. 2018;164:155–63. doi: 10.1016/j.neuroimage.2017.02.020.

12. Dumoulin SO, Fracasso A, van der Zwaag W, Siero JCW, Petridou N. Ultra-high field MRI: Advancing systems neuroscience towards mesoscopic human brain function. NeuroImage. 2018;168:345–57. doi: 10.1016/j.neuroimage.2017.01.028.

13. Huber L, Tse DHY, Wiggins CJ, Uludağ K, Kashyap S, Jangraw DC, Bandettini PA, Poser BA, Ivanov D. Ultra-high resolution blood volume fMRI and BOLD fMRI in humans at 9.4 T: Capabilities and challenges. NeuroImage. 2018;178:769–79. doi: 10.1016/j.neuroimage.2018.06.025.

14. Uğurbil K. Ultrahigh field and ultrahigh resolution fMRI. Current Opinion in Biomedical Engineering. 2021;18:100288. doi: 10.1016/j.cobme.2021.100288.

15. Scheffler K, Ehses P. High-resolution mapping of neuronal activation with balanced SSFP at 9.4 tesla. Magn Reson Med. 2016;76(1):163–71. Epub 2015/08/25. doi: 10.1002/mrm.25890. PubMed PMID: 26302451.

16. He Y, Wang M, Chen X, Pohmann R, Polimeni JR, Scheffler K, Rosen BR, Kleinfeld D, Yu X. Ultra-Slow Single-Vessel BOLD and CBV-Based fMRI Spatiotemporal Dynamics and Their Correlation with Neuronal Intracellular Calcium Signals. Neuron. 2018;97(4):925–39 e5. Epub 2018/02/06. doi: 10.1016/j.neuron.2018.01.025. PubMed PMID: 29398359; PMCID: PMC5845844.

17. He Y, Wang M, Yu X. High spatiotemporal vessel-specific hemodynamic mapping with multi-echo single-vessel fMRI. Journal of Cerebral Blood Flow & Metabolism. 2019;40(10):2098–114. doi: 10.1177/0271678X19886240.

18. Bolan PJ, Yacoub E, Garwood M, Ugurbil K, Harel N. In vivo micro-MRI of intracortical neurovasculature. Neuroimage. 2006;32(1):62–9. Epub 2006/05/06. doi: 10.1016/j.neuroimage.2006.03.027. PubMed PMID: 16675271.

19. Han S, Eun S, Cho H, Uludaǧ K, Kim S-G. Improved laminar specificity and sensitivity by combining SE and GE BOLD signals. NeuroImage. 2022;264:119675. doi: 10.1016/j.neuroimage.2022.119675.

20. Chen X, Jiang Y, Choi S, Pohmann R, Scheffler K, Kleinfeld D, Yu X. Assessment of single-vessel cerebral blood velocity by phase contrast fMRI. PLoS Biol. 2021;19(9):e3000923. Epub 2021/09/10. doi: 10.1371/journal.pbio.3000923. PubMed PMID: 34499636; PMCID: PMC8454982.

21. Scheffler K, Lehnhardt S. Principles and applications of balanced SSFP techniques. Eur Radiol. 2003;13(11):2409–18. Epub 2003/08/21. doi: 10.1007/s00330-003-1957-x. PubMed PMID: 12928954.

22. Miller KL. FMRI using balanced steady-state free precession (SSFP). Neuroimage. 2012;62(2):713–9. Epub 20111020. doi: 10.1016/j.neuroimage.2011.10.040. PubMed PMID: 22036996; PMCID: PMC3398389.

23. Yu X, Glen D, Wang S, Dodd S, Hirano Y, Saad Z, Reynolds R, Silva AC, Koretsky AP. Direct imaging of macrovascular and microvascular contributions to BOLD fMRI in layers IV-V of the rat whisker-barrel cortex. Neuroimage. 2012;59(2):1451–60. Epub 20110807. doi: 10.1016/j.neuroimage.2011.08.001. PubMed PMID: 21851857; PMCID: PMC3230765.

24. Drew PJ, Shih AY, Kleinfeld D. Fluctuating and sensory-induced vasodynamics in rodent cortex extend arteriole capacity. Proc Natl Acad Sci U S A. 2011;108(20):8473–8. Epub 2011/05/04. doi: 10.1073/pnas.1100428108. PubMed PMID: 21536897; PMCID: PMC3100929.

25. Silva AC, Koretsky AP, Duyn JH. Functional MRI impulse response for BOLD and CBV contrast in rat somatosensory cortex. Magn Reson Med. 2007;57(6):1110–8. Epub 2007/05/31. doi: 10.1002/mrm.21246. PubMed PMID: 17534912; PMCID: PMC4756432.

26. Yoshiyuki H, Bojana S, Afonso CS. Spatiotemporal Evolution of the Functional Magnetic Resonance Imaging Response to Ultrashort Stimuli. The Journal of Neuroscience. 2011;31(4):1440. doi: 10.1523/JNEUROSCI.3986-10.2011.

27. Uludag K, Muller-Bierl B, Ugurbil K. An integrative model for neuronal activity-induced signal changes for gradient and spin echo functional imaging. Neuroimage. 2009;48(1):150–65. Epub 20090527. doi: 10.1016/j.neuroimage.2009.05.051. PubMed PMID: 19481163.

28. Rabut C, Correia M, Finel V, Pezet S, Pernot M, Deffieux T, Tanter M. 4D functional ultrasound imaging of whole-brain activity in rodents. Nature Methods. 2019;16(10):994–7. doi: 10.1038/s41592-019-0572-y.

29. Chen Z, Zhou Q, Deán-Ben XL, Gezginer I, Ni R, Reiss M, Shoham S, Razansky D. Multimodal Noninvasive Functional Neurophotonic Imaging of Murine Brain-Wide Sensory Responses. Advanced Science. 2022;9(24):2105588. doi: 10.1002/advs.202105588.

30. Ovsepian SV, Jiang Y, Sardella TCP, Malekzadeh-Najafabadi J, Burton NC, Yu X, Ntziachristos V. Visualizing cortical response to optogenetic stimulation and sensory inputs using multispectral handheld optoacoustic imaging. Photoacoustics. 2020;17:100153. doi: 10.1016/j.pacs.2019.100153.

31. Duong TQ, Yacoub E, Adriany G, Hu X, Ugurbil K, Vaughan JT, Merkle H, Kim SG. High-resolution, spin-echo BOLD, and CBF fMRI at 4 and 7 T. Magn Reson Med. 2002;48(4):589–93. doi: 10.1002/mrm.10252. PubMed PMID: 12353274.

32. Larson PEZ, Gurney PT, Nishimura DG. Anisotropic Field-of-Views in Radial Imaging. IEEE Transactions on Medical Imaging. 2008;27(1):47–57. doi: 10.1109/TMI.2007.902799.

33. Feng L. Golden-angle radial MRI: basics, advances, and applications. Journal of Magnetic Resonance Imaging. 2022;56(1):45–62.

34. Winkelmann S, Schaeffter T, Koehler T, Eggers H, Doessel O. An optimal radial profile order based on the Golden Ratio for time-resolved MRI. IEEE Trans Med Imaging. 2007;26(1):68–76. Epub 2007/01/25. doi: 10.1109/TMI.2006.885337. PubMed PMID: 17243585.

35. Feng L, Grimm R, Block KT, Chandarana H, Kim S, Xu J, Axel L, Sodickson DK, Otazo R. Golden-angle radial sparse parallel MRI: combination of compressed sensing, parallel imaging, and golden-angle radial sampling for fast and flexible dynamic volumetric MRI. Magn Reson Med. 2014;72(3):707–17. Epub 2013/10/22. doi: 10.1002/mrm.24980. PubMed PMID: 24142845; PMCID: PMC3991777.

36. Toi PT, Jang HJ, Min K, Kim SP, Lee SK, Lee J, Kwag J, Park JY. In vivo direct imaging of neuronal activity at high temporospatial resolution. Science. 2022;378(6616):160–8. Epub 20221013. doi: 10.1126/science.abh4340. PubMed PMID: 36227975.

37. Yu X, Qian C, Chen DY, Dodd SJ, Koretsky AP. Deciphering laminar-specific neural inputs with line-scanning fMRI. Nat Methods. 2014;11(1):55–8. doi: 10.1038/nmeth.2730. PubMed PMID: 24240320; PMCID: PMC4276040.

38. Silva AC, Koretsky AP. Laminar specificity of functional MRI onset times during somatosensory stimulation in rat. Proceedings of the National Academy of Sciences. 2002;99(23):15182–7. doi: 10.1073/pnas.222561899.

39. Drew PJ, Mateo C, Turner KL, Yu X, Kleinfeld D. Ultra-slow Oscillations in fMRI and Resting-State Connectivity: Neuronal and Vascular Contributions and Technical Confounds. Neuron. 2020. Epub 2020/08/14. doi: 10.1016/j.neuron.2020.07.020. PubMed PMID: 32791040.

40. Pais-Roldán P, Biswal B, Scheffler K, Yu X. Identifying respiration-related aliasing artifacts in the rodent resting-state fMRI. Frontiers in Neuroscience. 2018;12:788.

41. Pais-Roldán P, Mateo C, Pan W-J, Acland B, Kleinfeld D, Snyder LH, Yu X, Keilholz S. Contribution of animal models toward understanding resting state functional connectivity. NeuroImage. 2021;245:118630. doi: 10.1016/j.neuroimage.2021.118630.

42. Robert T. How Much Cortex Can a Vein Drain? Downstream Dilution of Activation-Related Cerebral Blood Oxygenation Changes. NeuroImage. 2002;16(4):1062–7. doi: 10.1006/nimg.2002.1082.

43. Yacoub E, Duong TQ, Van De Moortele P-F, Lindquist M, Adriany G, Kim S-G, Uğurbil K, Hu X. Spin-echo fMRI in humans using high spatial resolutions and high magnetic fields. Magnet Reson Med. 2003;49(4):655–64. doi: 10.1002/mrm.10433.

44. Goense J, Merkle H, Logothetis NK. High-resolution fMRI reveals laminar differences in neurovascular coupling between positive and negative BOLD responses. Neuron. 2012;76(3):629–39. Epub 2012/11/13. doi: 10.1016/j.neuron.2012.09.019. PubMed PMID: 23141073; PMCID: PMC5234326.

45. Bianciardi M, Fukunaga M, van Gelderen P, de Zwart JA, Duyn JH. Negative BOLD-fMRI Signals in Large Cerebral Veins. Journal of Cerebral Blood Flow & Metabolism. 2010;31(2):401–12. doi: 10.1038/jcbfm.2010.164.

46. Kay K, Jamison KW, Zhang R-Y, Uğurbil K. A temporal decomposition method for identifying venous effects in task-based fMRI. Nature Methods. 2020;17(10):1033–9. doi: 10.1038/s41592-020-0941-6.

47. Caballero-Gaudes C, Reynolds RC. Methods for cleaning the BOLD fMRI signal. NeuroImage. 2017;154:128–49. doi: 10.1016/j.neuroimage.2016.12.018.

48. Polimeni JR, Lewis LD. Imaging faster neural dynamics with fast fMRI: A need for updated models of the hemodynamic response. Progress in Neurobiology. 2021;207:102174. doi: 10.1016/j.pneurobio.2021.102174.

49. Yu X, Glen D, Wang S, Dodd S, Hirano Y, Saad Z, Reynolds R, Silva AC, Koretsky AP. Direct imaging of macrovascular and microvascular contributions to BOLD fMRI in layers IV–V of the rat whisker–barrel cortex. NeuroImage. 2012;59(2):1451–60. doi: 10.1016/j.neuroimage.2011.08.001.

50. Tian P, Teng IC, May LD, Kurz R, Lu K, Scadeng M, Hillman EM, De Crespigny AJ, D’Arceuil HE, Mandeville JB, Marota JJ, Rosen BR, Liu TT, Boas DA, Buxton RB, Dale AM, Devor A. Cortical depth-specific microvascular dilation underlies laminar differences in blood oxygenation level-dependent functional MRI signal. Proc Natl Acad Sci U S A. 2010;107(34):15246–51. Epub 2010/08/11. doi: 10.1073/pnas.1006735107. PubMed PMID: 20696904; PMCID: PMC2930564.

51. Feng L, Axel L, Chandarana H, Block KT, Sodickson DK, Otazo R. XD-GRASP: golden-angle radial MRI with reconstruction of extra motion-state dimensions using compressed sensing. Magnet Reson Med. 2016;75(2):775–88.

