## Supplementary files for "High spatiotemporal resolution radial encoding single vessel fMRI"

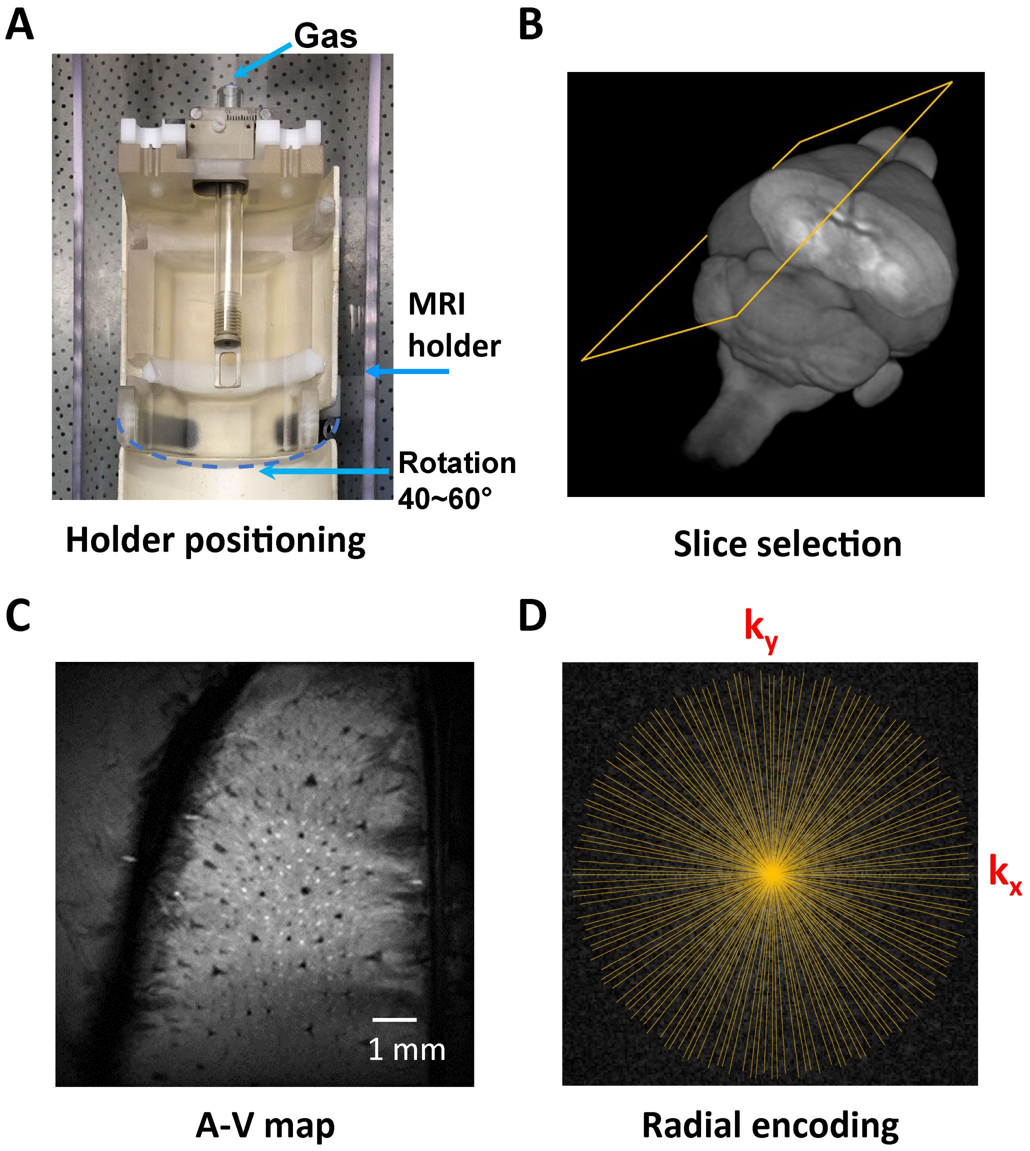


**Figure S1.** The 2D golden angle radial encoding MRI approach for *in vivo* fMRI experiment. (**A)** The rotatable animal holder for animal head positioning freedom in 14T MRI scanner. (**B)** 2D slice MRI selection for single-vessel radial encoding fMRI mapping of somatosensory cortex. (**C**) A representative anatomical 2D A-V (arteriole–venule) map acquired by multi-gradient echo (MGE) sequences. (**D)** The golden-angle radial encoding scheme. The radial encoding scheme allows single-vessel fMRI acquisition with an arbitrary number of profiles for the radial sampling of k-space.

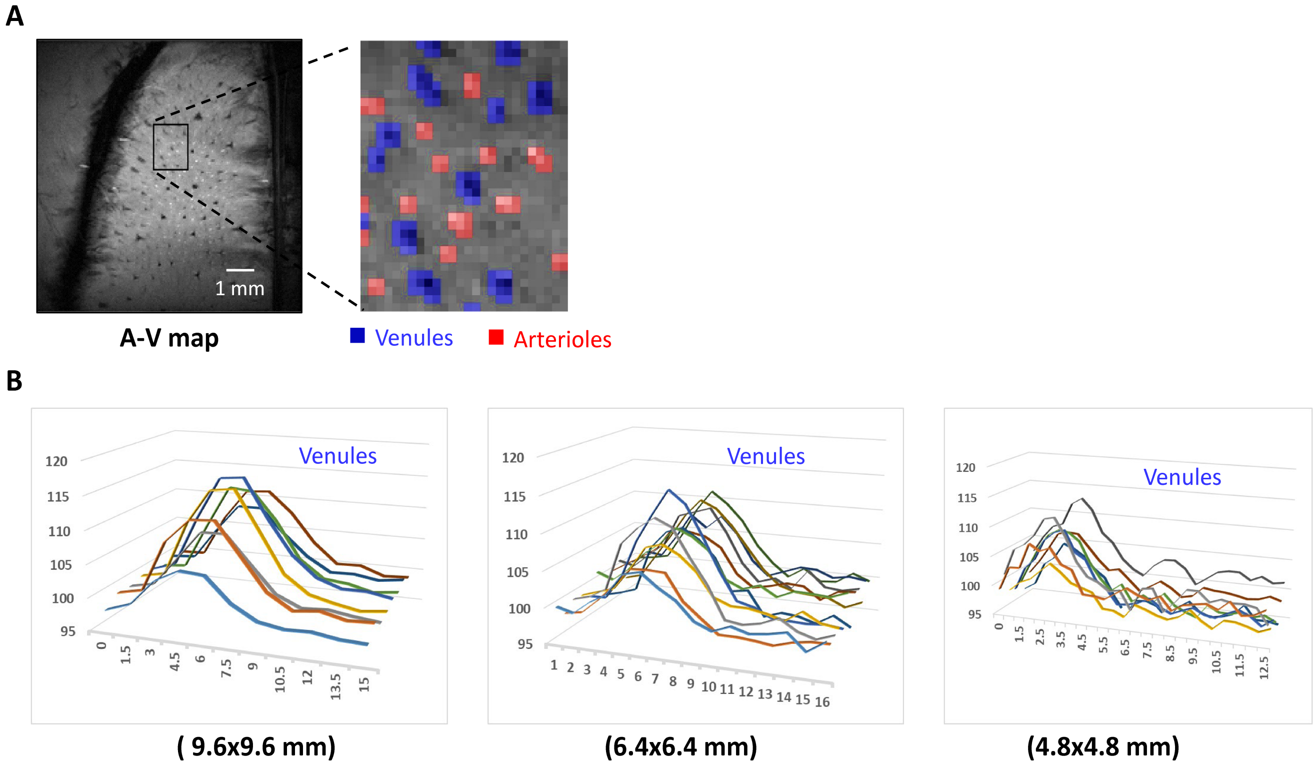


**Figure S2.** The representative venule-specific time course for radial encoding single vessel fMRI. (**A**) The individual arteriole and venule voxels were extracted from the A-V map with different signal intensities (venule voxels, blue; arteriole voxels, red). (**B**) Individual venule voxel signal at different FOV acquisitions (9.6x9.6, 6.4x6.4, and 4.8x4.8 mm$^2$) from one representative rat.

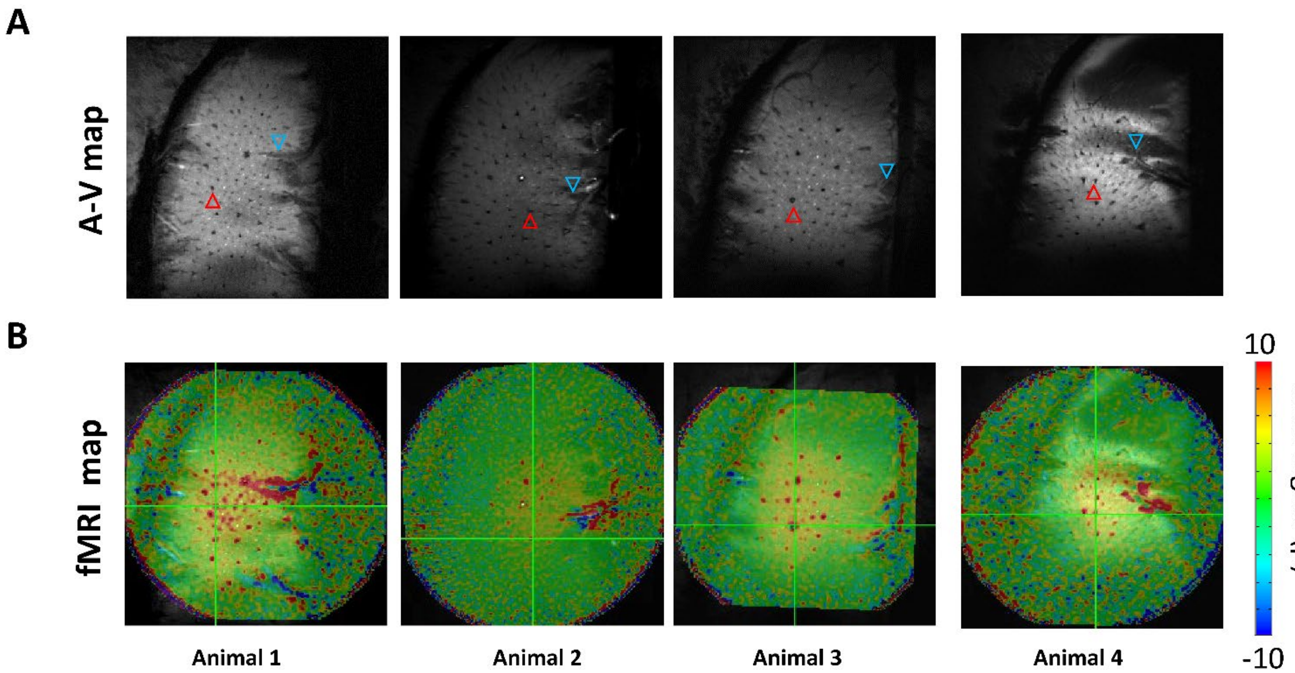


**Figure S3.** The positive BOLD signal and negative BOLD signals can be detected surrounding the pial veins with 100x100 μm resolution single-vessel fMRI. (**A**) 2-D slice A-V map of 4 representative animals. The red and blue triangle denotes the representative positive venules from the active cortex and negative BOLD signals surrounding the pial veins in the A-V map. (**B**) The corresponding fMRI map (100x100 μm resolution) was overlaid on the corresponding A-V map (4 animals).

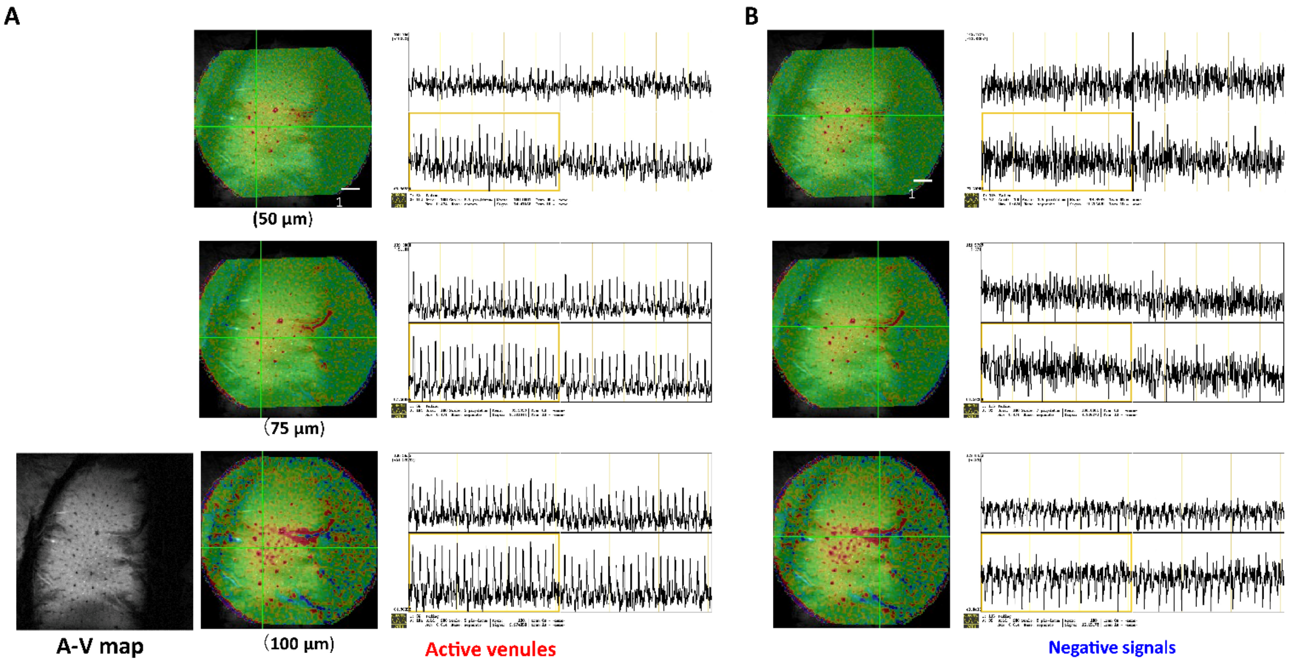


**Figure S4.** The fMRI mapping and time course of positive BOLD and negative BOLD signals at different resolution radial encoding based single-vessel fMRI. (**A**) Representative positive BOLD signal from one venule voxel from the active somatosensory cortex from different spatial resolutions (50, 75, and 100 µm). (**B**) The negative BOLD signal surrounding the pial veins was reduced with a higher spatial resolution single-vessel radial encoding fMRI.

### Supplementary Movie Legend

Supplementary movie 1.mp4

This movie shows the signal variations in radial encoding based single-vessel and bSSFP fMRI at the same resolution. The representative banding artifacts patterns evolved during the bSSFP acquisition due to the altered gradient/shimming coil temperature.
